# Simultaneous Determination of Contents of Flavonol Glycosides and Terpene Lactones in Ginkgo Biloba Tablets by Ultra High Performance Liquid Chromatography Tandem Single Quadrupole Mass Spectrometry Detector

**DOI:** 10.1101/2020.01.14.906347

**Authors:** Lin Ma, Jie Zhang, Wei-rong Jin, Siwang Wang

## Abstract

Ginkgo biloba leaf tablets is an effective ingredient in the treatment of cardiovascular and cerebrovascular diseases. In the process of drug production, the quality of ginkgo preparations is often controlled by measuring the content of seven ingredients in ginkgo leaves. To establish UPLC-MS multicomponent analysis method for ginkgo biloba tablets and to simultaneously determine the contents of quercetin (QUE), isorhamnetin(ISO), kaempferol(KAE) and GinkgolideA (GA),ginkgolideB(GB),ginkgolideC(GC) and bilobalide (BB) in ginkgo tablets. Waters Xbridge C18(4.6×150mm,3.5um) column was used, mobile phase A was acetonitrile and mobile phase B was water (containing 0.10% formic acid). The injection volume was 10μL.Negative ion mode monitoring was conducted with ESI. Scanning range:m/z100∼1400.The detection ions of the seven tested components includem/z301.0(QUE),m/z284.9(KAE),m/z315.1(ISO),m/z453.1(GA),m/z423.1(G B),m/z439.0(GC)and m/z325.0(BB), respectively. Within a space of 10min, flavonoids and terpene lactones in ginkgo biloba tablets were completely separated. The peak area exhibited an excellent linear relationship with the concentration. The sample recovery rate ranged from 91.74% to 109.77%.Precision RSDs of within-day and between-day were lower than 2.879% and 3.928% respectively. The method for determination of seven components in ginkgo biloba tablets displays good repeatability, recovery rate and precision, for which it can be applied to quality control of ginkgo biloba tablets.

## 1. Introduction

The major component of ginkgo biloba tablets is ginkgo extract, the main chemical components of which include ginkgo flavone and ginkgolide. In traditional Chinese medicine, it is known that Ginkgo biloba leaves and ginkgo nuts taste sweet, bitter and astringent. It is mild, with its leaves capable of the effects to promote blood circulation, nourish heart, as well as astringe lungs and intestine. The kernel of its seeds has the effects of moistening lung, alleviating asthma, reducing cough, inducing diuresis, preventing white ooze, inhibiting worms, relieving hangover, etc. Ginkgo biloba exocarp is sweet in taste, mild in nature and has the effects of enhancing vigour and tonifying deficiency [1-8]. Ginkgo biloba flavonoids are primarily present in ginkgo biloba leaves and seed kernels, with an especially high content found in ginkgo biloba leaves. Among them, quercetin, isorhamnetin and ft-nai have higher contents and are the main components of ginkgo biloba flavonoids. In the process of drug production, the quality of ginkgo biloba preparations is often controlled by detecting the contents of these three flavonoid aglycones [2-5]. Ginkgolide is an extraordinary component of ginkgo biloba and is contained in the seeds, leaves, roots and stems of ginkgo biloba. Ginkgolide A, B, C and bilobalide are also considered to be significant indicators of quality control for ginkgo biloba preparations [2-7].

At present, there are a variety of different methods for detecting ginkgo flavone and ginkgolide during ginkgo preparations [9-13]. However, there are few reports focused on multi-component mixed analysis of ginkgo preparations and there remain no reports on simultaneous determination and research into the seven components of ginkgo flavone and ginkgolide in ginkgo biloba tablets. In terms of detection instruments, ginkgolides are not absorbed in the ultraviolet region, for which it cannot be determined using HPLC-UV method. Nowadays HPLC-ELSD method is widely used [14,15]. However, the evaporative light detector has drawbacks of higher noise, lower sensitivity and poor stability. It requires complex sample processing to be used for detection [16]. In this experiment, simultaneous qualitative and quantitative analysis was conducted of ginkgo flavone and ginkgolide compounds without ultraviolet absorption by Ultra High Performance Liquid Chromatography Tandem Single Quadr-upole Mass Spectrometry Detector(UPLC-MS).

## 2. Results

### 2.1 Quality Evaluation of Total Flavonoid Glycosides and Terpene Lactones Contents in Ginkgo Biloba Tablets

#### 2.1.1 Determination of Total Flavonoid Glycosides Contents in Ginkgo Biloba Tablets

As specified in the first part of the 2015 edition of the Chinese Pharmacopoeia [17],the total flavonol glycosides in specification A shall be 19.2mg·tablet^-1^ as a minimum and the total flavonol glycosides content in specification B shall be 9.6mg·tablet^-1^ as a minimum (n=3). The results demonstrated that the content of total flavonol glycosides in ten batches of ginkgo bilobo tablets produced by five manufacturers was compliant,as shown in Table1.

**Table 1.** Determination of total flavonoid glycosides contents in ginkgo biloba tablets (n=3).

| Numbering | sample | Specification (mg) | Mass fraction ($\text{mg}\cdot\text{tablet}^{-1}$) |
| --- | --- | --- | --- |
| 1 | A1 | 19.2 | 23.1 |
| 2 | A2 | 19.2 | 22.2 |
| 3 | A3 | 19.2 | 22.4 |
| 4 | A4 | 19.2 | 22.3 |
| 5 | A5 | 19.2 | 22.0 |
| 6 | A6 | 19.2 | 22.9 |
| 7 | B1 | 9.6 | 13.5 |
| 8 | B2 | 9.6 | 12.6 |
| 9 | B3 | 9.6 | 11.3 |
| 10 | B4 | 9.6 | 12.1 |

#### 2.1.2 Determination of Terpene Lactones Contents in Ginkgo Biloba Tablets

As specified in the first part of the 2015 edition of the Chinese Pharmacopoeia [17], the terpene lactones in specification A shall be 4.8mg·tablet^-1^ as a minimum and the terpene lactones content in specification B shall be 2.4mg·tablet^-1^ as a minimum(n=3). The results indicated that the content of terpene lactones in ten batches of ginkgo bilobo tablets produced by five manufacturers was compliant, as shown in Table 2.

**Table 2.** Determination of terpene lactones contents in ginkgo biloba tablets (n=3).

| Numbering | sample | Specifications (mg) | Mass fraction ($\text{mg}\cdot\text{tablet}^{-1}$) |
| --- | --- | --- | --- |
| 1 | A1 | 4.8 | 5.4 |
| 2 | A2 | 4.8 | 5.1 |
| 3 | A3 | 4.8 | 6.0 |
| 4 | A4 | 4.8 | 5.0 |
| 5 | A5 | 4.8 | 5.9 |
| 6 | A6 | 4.8 | 9.0 |
| 7 | B1 | 2.4 | 3.1 |
| 8 | B2 | 2.4 | 3.1 |
| 9 | B3 | 2.4 | 3.4 |
| 10 | B4 | 2.4 | 3.1 |

### 2.2 Simultaneous Determination of 7 Components in Ginkgo Biloba Tablets by UPLC-MS and Validation of Methodology

#### 2.2.1 Standard Curve, Detection Limit and Quantitative Limit

A precise measurement was taken of the series of standard solutions and the determination was performed according to the “chromatographic conditions”(n=3). The concentrations of QUE, KAE, ISO, GA, GB, GC, BB were taken as abscissa, the peak areas were taken as ordinate and a linear regression equation was derived. The results indicated that the linear relationship of each standard was excellent within a certain range. The mixed reference substance solution was diluted incrementally. The concentration of each reference substance when S/N=10:1 and S/N=3:1 was regarded as the quantitative limit and detection limit [18]. The quantitative limit range was 5.0*10^−6^ to 1.0*10^-5^, mg·mL^-1^ and the detection limit range was 1.0*10^-5^ to 2.5*10^-5^ mg·mL^-1^, as shown in Table 3.

**Table 3.** Standard curve of 7 components in ginkgo biloba tablets (n=3).

| Components | Linear equation | $r^2$ | Concentration range<br>($\text{mg}\cdot\text{mL}^{-1}$) | LOQ<br>($\text{mg}\cdot\text{mL}^{-1}$) | LOD<br>($\text{mg}\cdot\text{mL}^{-1}$) |
| --- | --- | --- | --- | --- | --- |
| QUE | $y=21731+11.91x$ | 0.9997 | 0.005~0.100 | $1.0\times 10^{-5}$ | $2.5\times 10^{-5}$ |
| KAE | $y=119304+14.53x$ | 0.9991 | 0.005~0.100 | $1.0\times 10^{-5}$ | $2.5\times 10^{-5}$ |
| ISO | $y=102424+16.15x$ | 0.9992 | 0.005~0.100 | $1.0\times 10^{-5}$ | $2.5\times 10^{-5}$ |
| GA | $y=44533+11.65x$ | 0.9992 | 0.005~0.100 | $5.0\times 10^{-6}$ | $1.0\times 10^{-5}$ |
| GB | $y=66218+12.03x$ | 0.9997 | 0.005~0.100 | $5.0\times 10^{-6}$ | $1.0\times 10^{-5}$ |
| GC | $y=42527+8.44x$ | 0.9996 | 0.005~0.100 | $5.0\times 10^{-6}$ | $1.0\times 10^{-5}$ |
| BB | $y=61237+12.66x$ | 0.9997 | 0.005~0.100 | $5.0\times 10^{-6}$ | $1.0\times 10^{-5}$ |

#### 2.2.2 Sample Recovery Rate

After one tablet of sample A1 was taken and re-extracted.1000μL of filtered extract was taken. According to the ratio of 1:0.5(low),1:1.0 (medium) and 1:1.5(high) of the content of each measured chemical substance, standard solution with various substance concentrations of 1mg·mL^-1^ was added respectively. In addition, the recovery rates of high, medium and low concentration substances were tested after the standard was added. Based on the determination of “chromatographic conditions”(n=3), the recovery rate of this method was 91.74%∼109.77%. The results confirmed that the recovery rate of this method could satisfy the requirements, as shown in Table 4.

**Table 4.** The recovery rate of seven components in ginkgo biloba tablets (n=3).

| Component | Sample content | Addition of standard substance | Measured quantity | Recovery rate | Average recovery rate | RSD |
| --- | --- | --- | --- | --- | --- | --- |
|  | (mg) | (mg) | (mg) | (%) | (%) | (%) |
| QUE | 0.112 | 0.0560 | 0.1735 | 109.82 | 101.29 | 2.19 |
|  |  | 0.1120 | 0.2185 | 95.08 |  | 0.41 |
|  |  | 0.1680 | 0.2783 | 98.97 |  | 3.00 |
| KAE | 0.213 | 0.1065 | 0.3258 | 105.95 | 97.65 | 1.16 |
|  |  | 0.2130 | 0.4131 | 93.92 |  | 0.87 |
|  |  | 0.3195 | 0.5104 | 93.09 |  | 1.54 |
| ISO | 0.175 | 0.0875 | 0.2707 | 109.42 | 102.04 | 3.14 |
|  |  | 0.1750 | 0.3587 | 104.97 |  | 3.15 |
|  |  | 0.2625 | 0.4158 | 91.74 |  | 0.00 |
| GA | 0.069 | 0.0345 | 0.1056 | 106.03 | 101.93 | 1.65 |
|  |  | 0.0690 | 0.1389 | 101.25 |  | 0.20 |
|  |  | 0.1035 | 0.1710 | 98.51 |  | 3.23 |
| GB | 0.046 | 0.0230 | 0.0694 | 101.82 | 101.18 | 2.52 |
|  |  | 0.0460 | 0.0920 | 99.94 |  | 2.97 |
|  |  | 0.0690 | 0.1162 | 101.78 |  | 2.79 |
| GC | 0.066 | 0.0330 | 0.1007 | 105.28 | 104.28 | 2.37 |
|  |  | 0.0660 | 0.1357 | 105.62 |  | 0.41 |
|  |  | 0.0990 | 0.1669 | 101.96 |  | 2.39 |
| BB | 0.125 | 0.0625 | 0.1935 | 109.58 | 99.19 | 1.71 |
|  |  | 0.1250 | 0.2415 | 93.17 |  | 0.97 |
|  |  | 0.1875 | 0.3028 | 94.84 |  | 0.79 |

#### 2.2.3 Precision

A precise measurement was taken of the same mixed reference substance solution. The mixed standard solution of low, medium and high concentration was prepared and determined based on “chromatographic conditions” (n=6). Intra-day precision refers to six parallel tests within one day. Daytime precision refers to two parallel tests within one day for three consecutive days. The results confirmed that the intra-day and inter-day precision of the method could satisfy the requirements, as shown in Table 5.

**Table 5.**
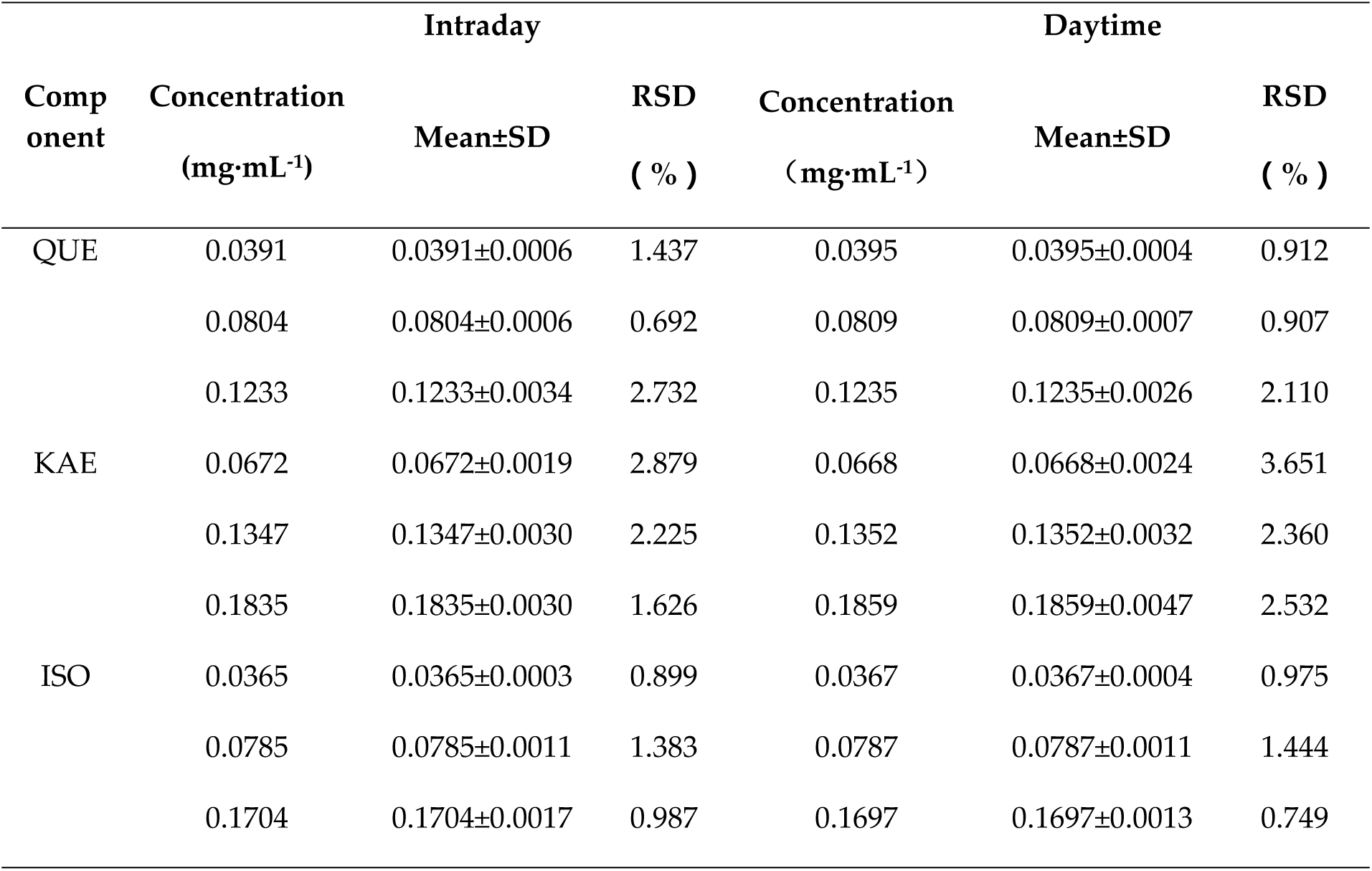
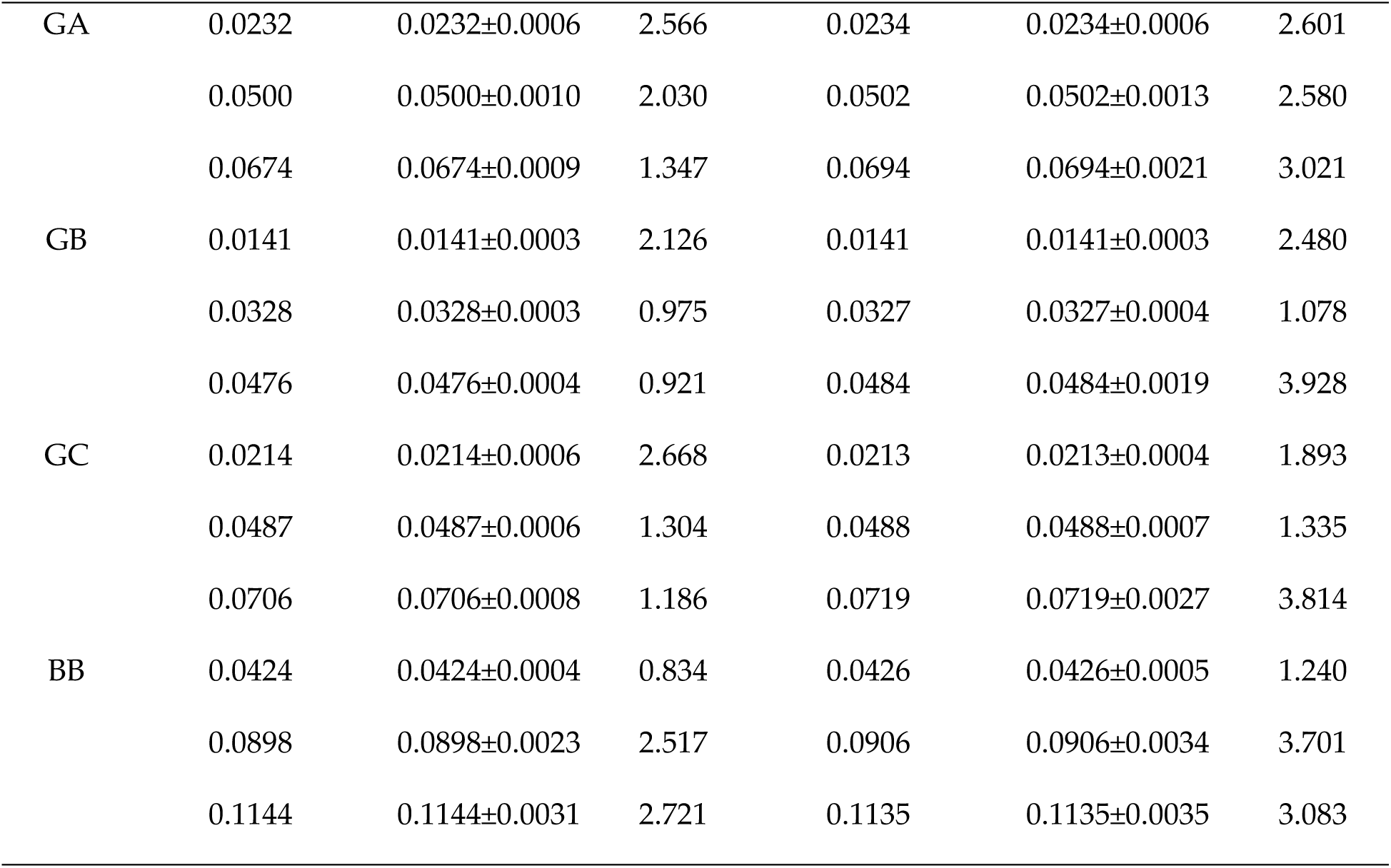
Intraday and daytime precision of seven components in ginkgo biloba tablets (n=6, 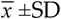).

#### 2.2.4 Repeatable

Sample 1 was taken. Five sample solutions were prepared in parallel according to the preparation items of the sample solution and determined based on the “chromatographic conditions”(n=3). The results indicated that the method was capable of excellent repeatability, as shown in Table 6.

**Table 6.** Repeatability of seven components in ginkgo biloba tablets (n=3).

| Sample | Measured content (mg·mL<sup>-1</sup> ) |  |  |  |  | Average | RDS |
| --- | --- | --- | --- | --- | --- | --- | --- |
| Component | 1 | 2 | 3 | 4 | 5 | (mg·mL<sup>-1</sup> ) | (%) |
| QUE | 0.1232 | 0.1226 | 0.1231 | 0.1231 | 0.1236 | 0.1231±0.00030 | 0.27 |
| KAE | 0.1945 | 0.1956 | 0.1943 | 0.1935 | 0.1959 | 0.1948±0.00154 | 0.79 |
| ISO | 0.1694 | 0.1721 | 0.1707 | 0.1726 | 0.1706 | 0.1711±0.00128 | 0.74 |
| GA | 0.0647 | 0.0668 | 0.0663 | 0.0669 | 0.0662 | 0.0661±0.00046 | 0.69 |
| GB | 0.0481 | 0.0486 | 0.0487 | 0.0487 | 0.0487 | 0.0486±0.00018 | 0.35 |
| GC | 0.0705 | 0.0720 | 0.0721 | 0.0727 | 0.0721 | 0.0719±0.00034 | 0.45 |
| BB | 0.1115 | 0.1115 | 0.1111 | 0.1112 | 0.1110 | 0.1113±0.00068 | 0.61 |

#### 2.2.5 Stability

Sample 1 was taken. According to the preparation items of the sample solution, seven samples of the sample solution were prepared in parallel. Based on the “chromatographic conditions”(n=3), the samples were injected for detection at 0, 4, 8, 12, 18, 24 and 48 hours respectively. The results revealed that the sample solution was capable of remarkable stability within forty-eight hours, as shown in Table 7.

**Table 7.**
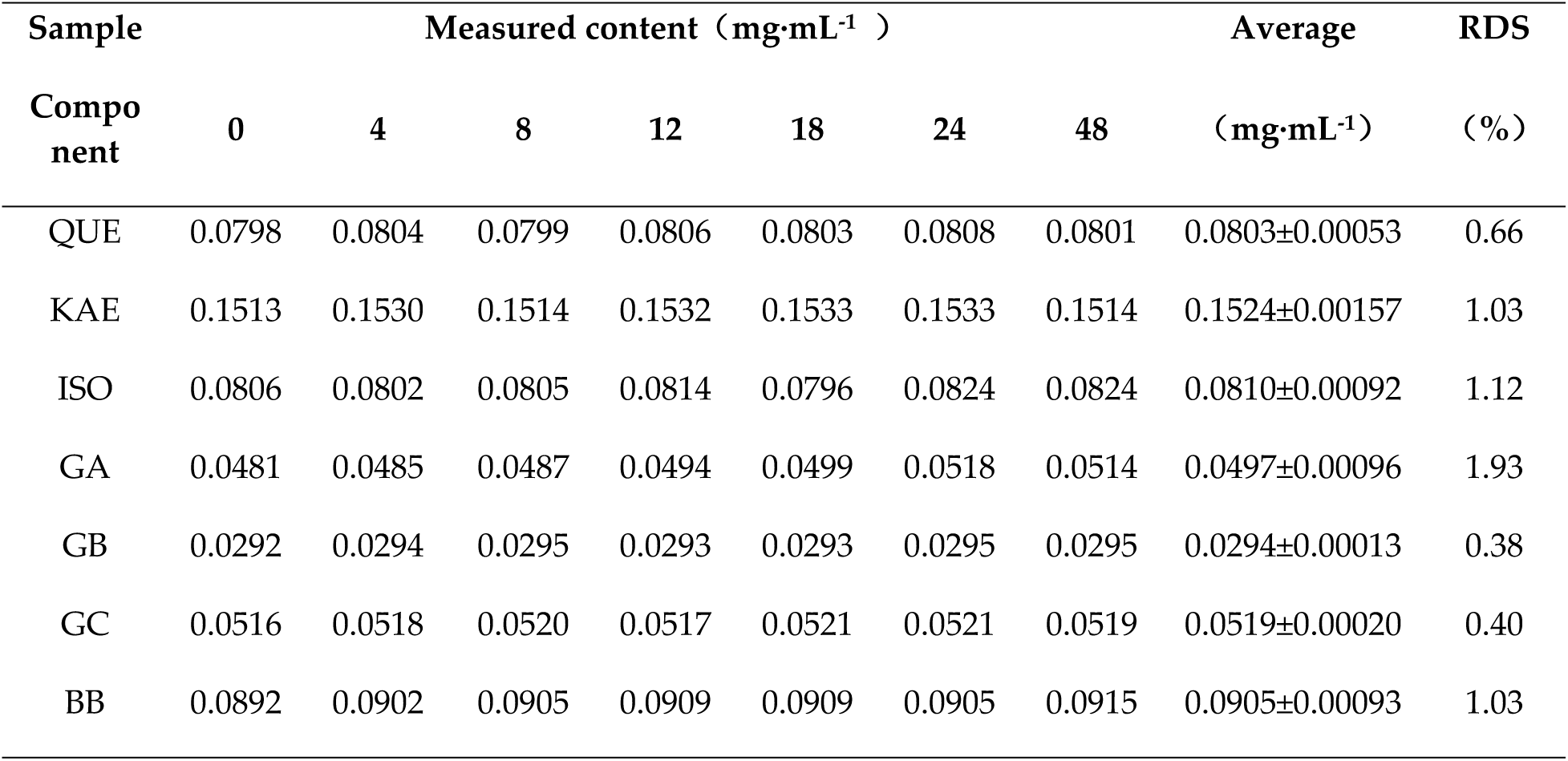
Stability of seven components in ginkgo biloba tablets (n=3).

| Sample | Measured content (mg·mL<sup>-1</sup>) |  |  |  |  |  |  | Average | RDS |
| --- | --- | --- | --- | --- | --- | --- | --- | --- | --- |
| Component | 0 | 4 | 8 | 12 | 18 | 24 | 48 | (mg·mL<sup>-1</sup>) | (%) |
| QUE | 0.0798 | 0.0804 | 0.0799 | 0.0806 | 0.0803 | 0.0808 | 0.0801 | 0.0803±0.00053 | 0.66 |
| KAE | 0.1513 | 0.1530 | 0.1514 | 0.1532 | 0.1533 | 0.1533 | 0.1514 | 0.1524±0.00157 | 1.03 |
| ISO | 0.0806 | 0.0802 | 0.0805 | 0.0814 | 0.0796 | 0.0824 | 0.0824 | 0.0810±0.00092 | 1.12 |
| GA | 0.0481 | 0.0485 | 0.0487 | 0.0494 | 0.0499 | 0.0518 | 0.0514 | 0.0497±0.00096 | 1.93 |
| GB | 0.0292 | 0.0294 | 0.0295 | 0.0293 | 0.0293 | 0.0295 | 0.0295 | 0.0294±0.00013 | 0.38 |
| GC | 0.0516 | 0.0518 | 0.0520 | 0.0517 | 0.0521 | 0.0521 | 0.0519 | 0.0519±0.00020 | 0.40 |
| BB | 0.0892 | 0.0902 | 0.0905 | 0.0909 | 0.0909 | 0.0905 | 0.0915 | 0.0905±0.00093 | 1.03 |

#### 2.2.6 Simultaneous Determination of Seven Components Content in Ginkgo Biloba Tablets

Totally ten batches of ginkgo leaf samples were taken and determined based on the terms of “preparation of test solution” and “chromatographic conditions”(n=3). The established analysis method was effective in separating the components in ginkgo biloba tablets and a quantitative analysis was performed of seven components. After calculation, the results revealed that QUE, KAE, ISO and BB are higher than other components in ten batches of ginkgo biloba tablets, as shown in Table 8. The LC-MS spectra of the 7 components reference substance were observed to be consistent with the corresponding component spectra of the sample, as shown in Figures 1 and 2. And the primary mass spectra of the reference substance and the sample are presented in Figures 3, 4, 5, 6, 7, 8 and 9.

**Table 8.** Contents of seven components in ginkgo biloba tablets in different batches (n=3, 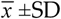, mg·g^-1^).

| Sample |  |  |  |  |  |  |  |
| --- | --- | --- | --- | --- | --- | --- | --- |
| /Component | QUE | KAE | ISO | GA | GB | GC | BB |
| A1 | 5.363±0.007 | 10.202±0.051 | 8.357±0.014 | 3.309±0.005 | 2.214±0.007 | 3.139± 0.004 | 5.989±0.001 |
| A2 | 5.684±0.002 | 10.128±0.012 | 8.291±0.007 | 3.387±0.003 | 2.392±0.004 | 3.199± 0.004 | 6.356±0.002 |
| A3 | 4.907±0.005 | 7.749±0.012 | 6.753±0.006 | 2.632±0.020 | 1.904±0.004 | 2.803±0.007 | 4.496±0.007 |
| A4 | 5.826±0.003 | 7.940±0.027 | 7.933±0.002 | 2.941±0.002 | 2.015±0.007 | 2.798± 0.002 | 4.847±0.008 |
| A5 | 10.658±0.007 | 13.680±0.009 | 14.492±0.006 | 4.240±0.002 | 3.138±0.030 | 4.274±0.011 | 6.454±0.002 |
| A6 | 5.493±0.006 | 7.980±0.006 | 7.245±0.002 | 3.222±0.002 | 1.896±0.001 | 2.405±0.013 | 4.334±0.004 |
| B1 | 4.306±0.003 | 4.793±0.002 | 4.381±0.003 | 1.767±0.014 | 1.247±0.004 | 1.646±0.005 | 3.332±0.001 |
| B2 | 4.342±0.005 | 5.168±0.005 | 4.948±0.004 | 2.001±0.002 | 1.368±0.001 | 1.846±0.006 | 3.504±0.005 |
| B3 | 5.634±0.004 | 7.157±0.024 | 6.649±0.025 | 2.754±0.004 | 1.736±0.002 | 1.953±0.002 | 4.053±0.002 |
| B4 | 5.318±0.004 | 7.100±0.007 | 6.744±0.026 | 2.606±0.006 | 1.646±0.010 | 1.841±0.047 | 3.731±0.004 |

**Figure 1.**
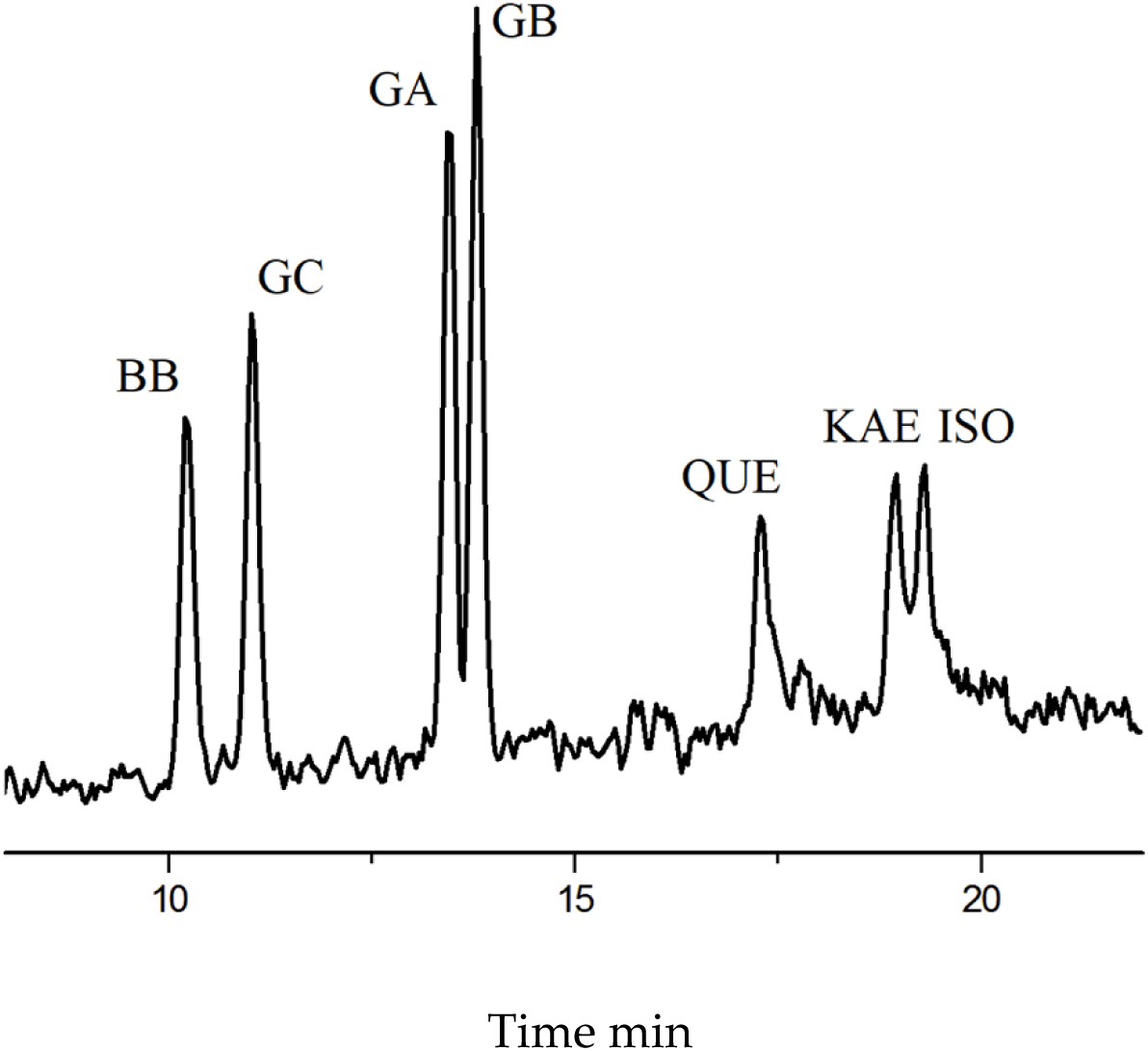
LC-MS chromatogram of seven components reference substance from ginkgo biloba.

**Figure 2.**
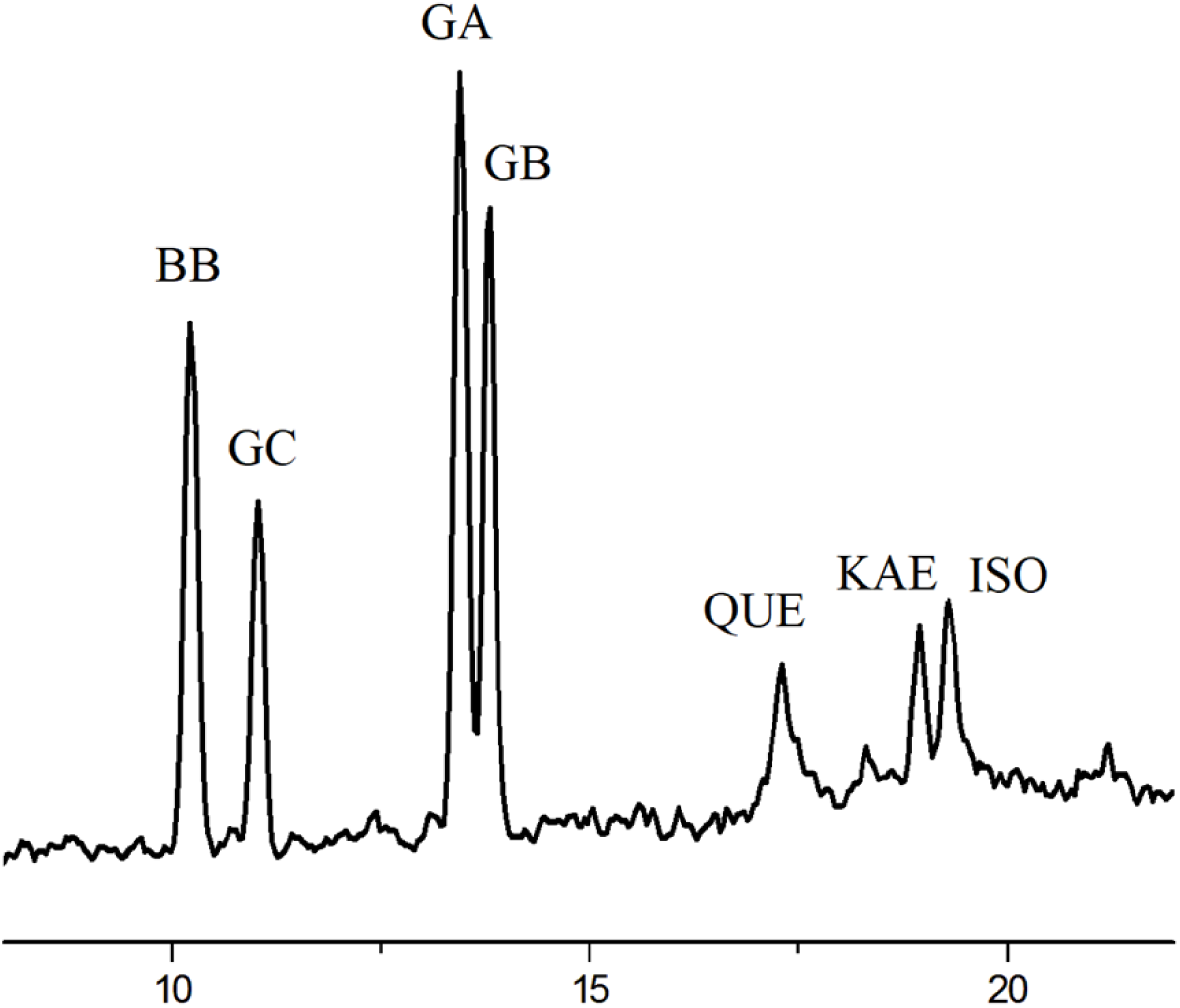
LC-MS chromatogram of seven component samples from ginkgo biloba tablets.

**Figure 3.**
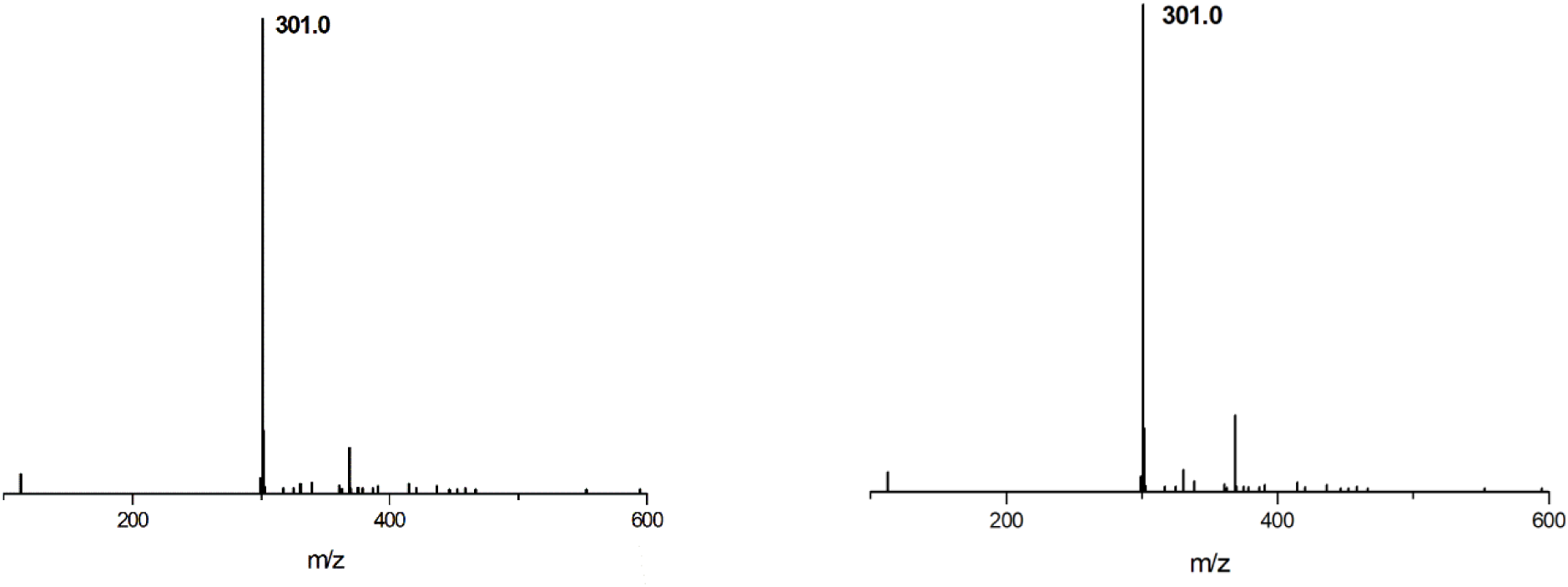
Primary mass spectrum of QUE reference substance (left) and sample (right).

**Figure 4.**
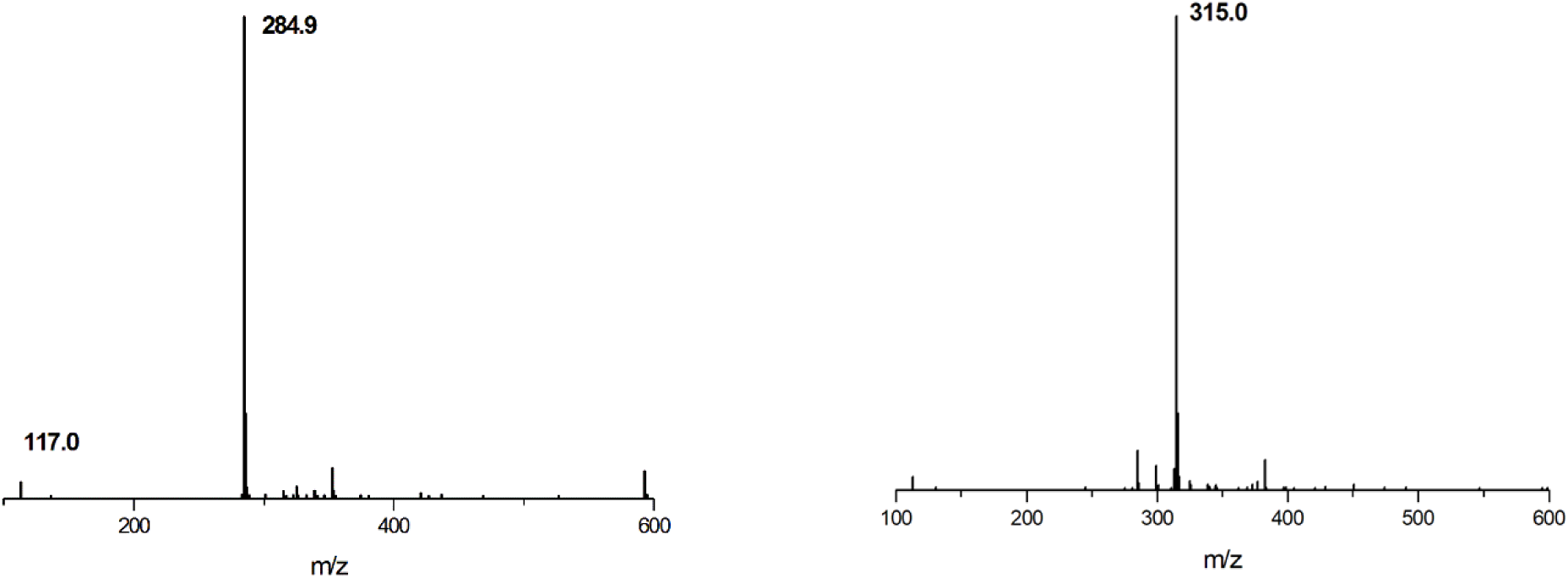
Primary mass spectrum of KAE reference substance (left) and sample (right).

**Figure 5.**
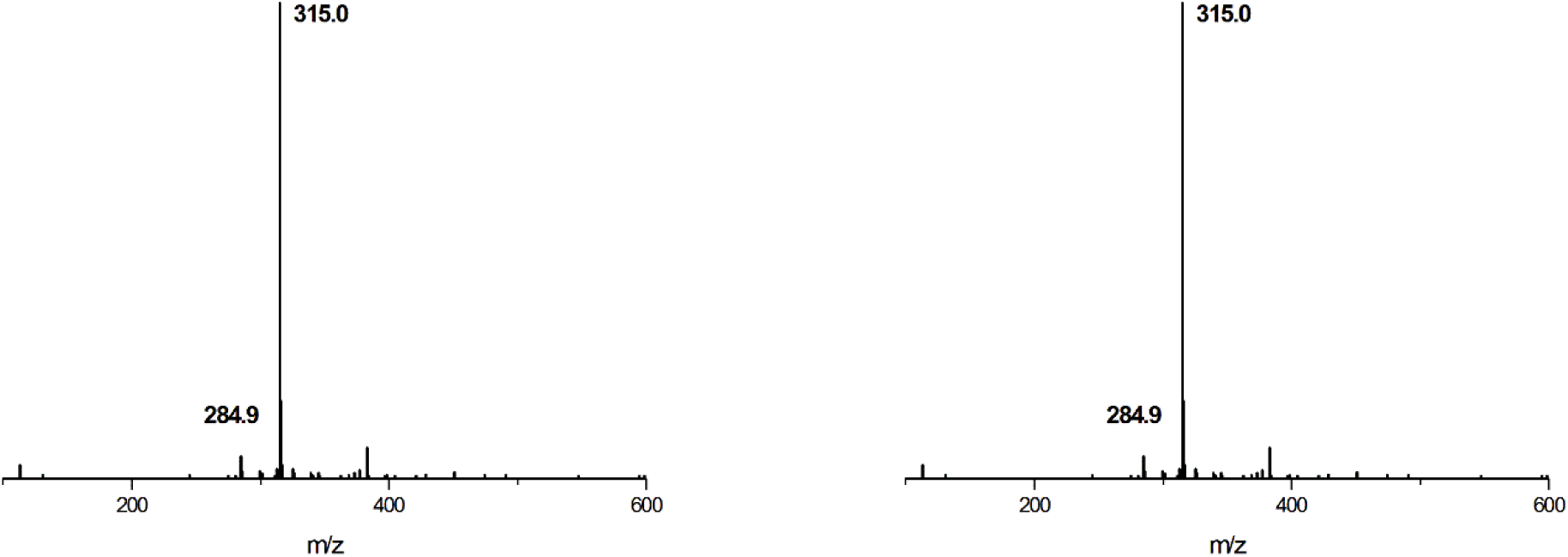
Primary mass spectrum of ISO reference substance (left) and sample (right).

**Figure 6.**
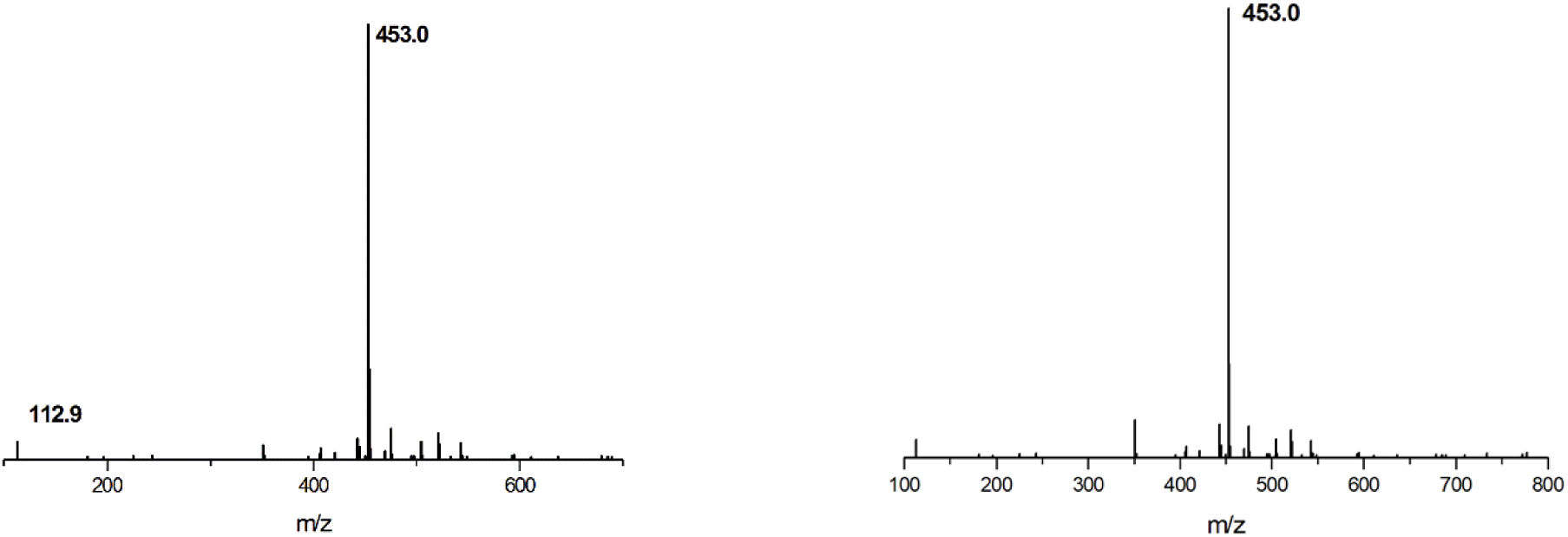
Primary mass spectrum of GA reference substance (left) and sample (right).

**Figure 7.**
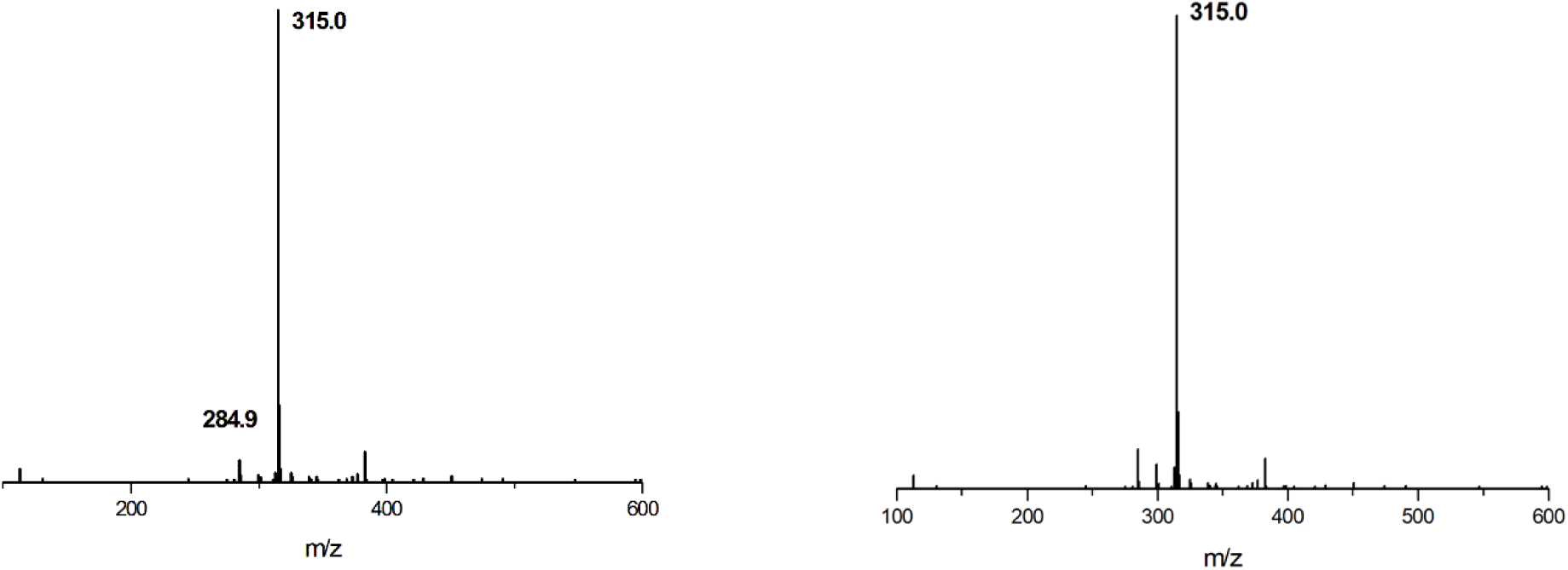
Primary mass spectrum of GB reference substance (left) and sample (right).

**Figure 8.**
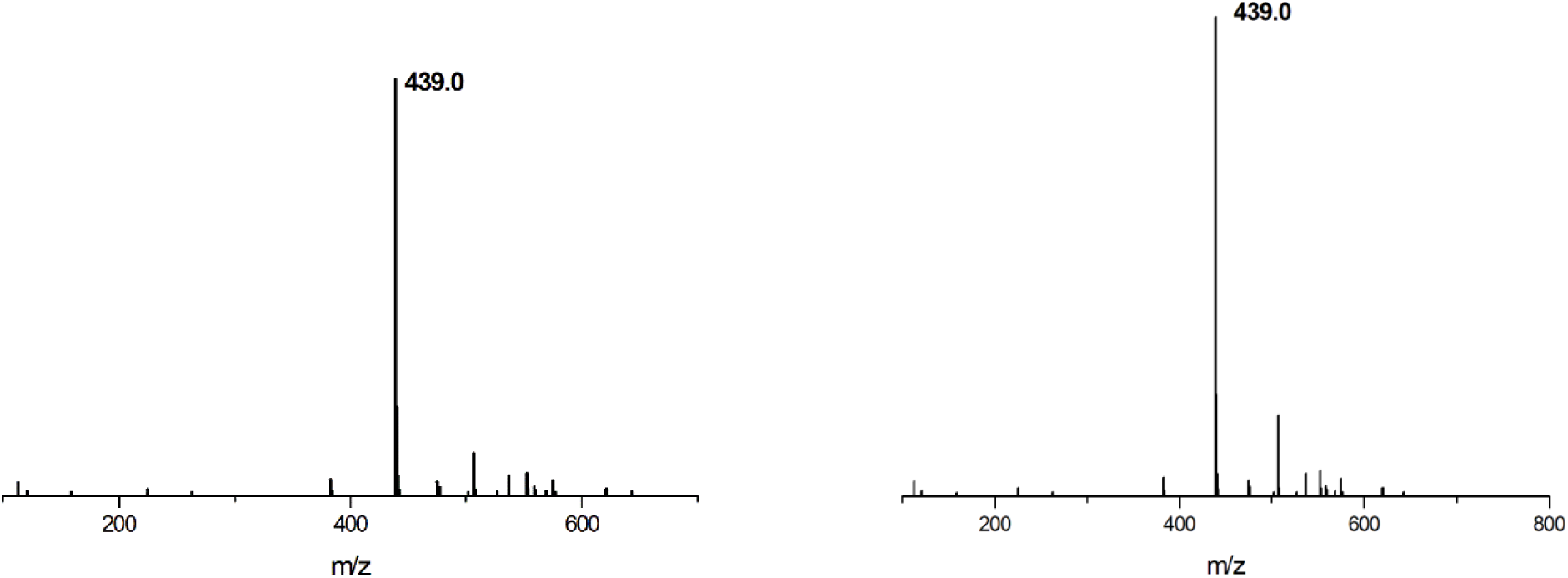
Primary mass spectra of GC reference substance (left) and sample (right).

**Figure 9.**
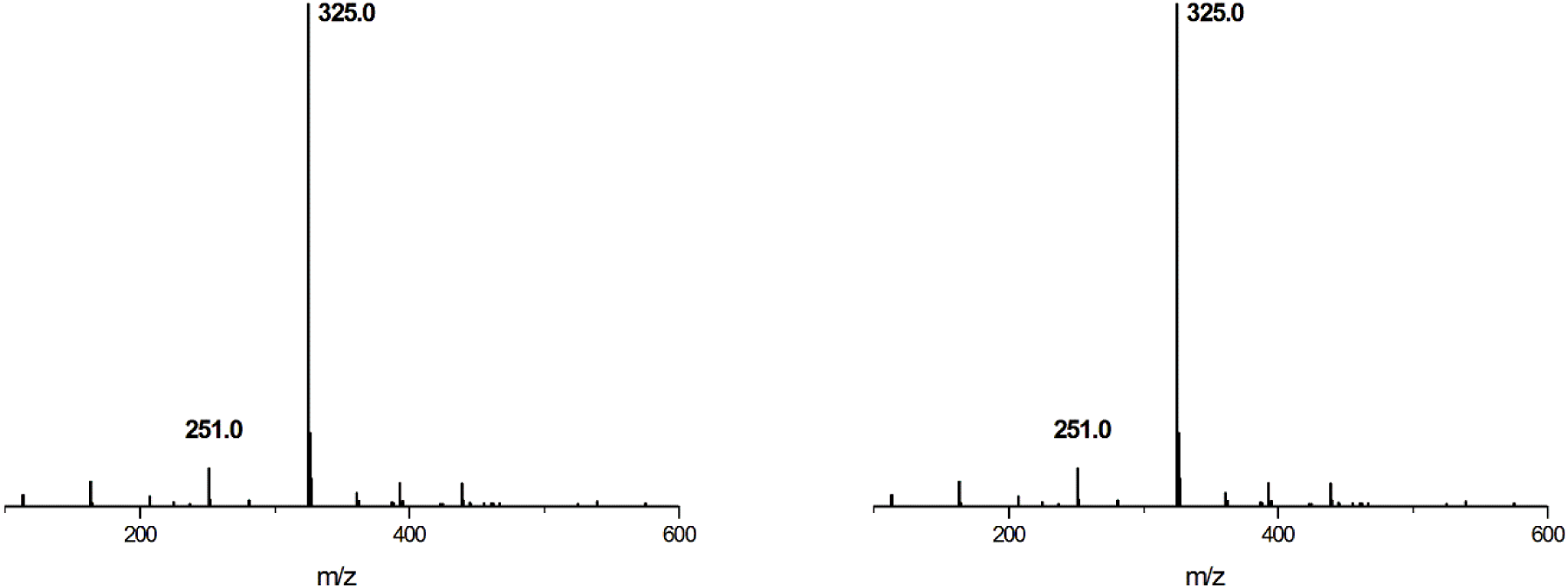
Primary mass spectrum of BB reference substance (left) and sample (right).

## 3. Discussion

High performance liquid chromatography-mass spectrometry (HPLC-MS) has been extensively applied to the quality research of drugs due to its impressive specificity and higher sensitivity [19-24]. In this experiment, the greatest advantage displayed by the UPLC-MS was simple operation. It took as little as five minutes of mass spectrometry equilibrium time to complete qualitative analysis of samples. Moreover, it can be used for qualitative and quantitative analysis of compounds that is incapable of ultraviolet absorption. Negativeion mode monitoring was conducted by Electron Spray Ionization and the results demonstrated that the relative molecular mass parameters were respectively QUE301.0,KAE284.9, ISO315.1,GA453.1,GB423.1,GC439.0 and BB325.0, which was discovered to be consistent with literature reports [22].Not only does this method provide reference and basis for the quality control of ginkgo biloba tablets, it also lays a solid foundation for further study on the pharmacodynamical characteristics of in vivo index components of ginkgo biloba after oral administration in rats. However, this method remains subjected to various limitations and some drawbacks are exposed. In this sense, further research is deemed necessary to improve the quality control of ginkgo biloba tablets and to figure out their efficacy.

## 4. Materials and Methods

### 4.1 Instruments and Drug Testing

#### Measuring apparatus

Electronic Balance, METTLER TOLEDO, USA; Ultrasonic Instrument,

shanghai Yixin,China;Centrifuge, Zhongke,China;Ultra high performance liquid chromatography system (UHPLC), Agilent 1260;Mass spectrometer (MS), Agilent 6120;N-EVAP -24 Organomation of the United States.

#### Standard substance

quercetin (QCT) batch number 181123, content ≥98.05%, kaempferol (KAE) batch number 10128, content ≥98.54%, isorhamnetin (ISR) batch number 181217, content ≥98.07%, ginkgolide A(GA) batch number 180412, content ≥99.32%, ginkgolideB(GB) batch number 180210, content ≥98.97%, ginkgolide C(GC) batch number 180330, content ≥98.8%, bilobalide (BB) batch number 180615, content ≥99.69%. They are all purchased from Beijing Century Aoke Biotechnology Co.LTD.

#### Reference substance

ginkgo biloba tablets five manufacturers,ten batch numbers,specifications:

19.2/4.8mg·tablet-1and 9.6/2.4mg·tablet^-1^.They are designated A1, A2, A3, A4, A5, A6, B1, B2, B3 and B4 respectively.

#### Reagent

methanol,chromatographicall pure,Thermo Fisher of USA; Acetonitrile, Chromatographically pure,Thermofisher of USA;Dichloromethane, Analytical pure, national medicine reagent chemical reagent co.LTD;Deionized water,self-made in laboratory, other reagents are all analytically pure.

### 4.2 content determination

#### 4.2.1 Quality Evaluation of Total Flavonoid Glycosides and Terpene Lactones in Ginkgo Biloba Tablets

According to the content determination method of total flavonol glycosides and terpenoid lactones in ginkgo biloba tablets of the first part of Chinese pharmacopoeia 2015 edition [17],ten batches of ginkgo biloba tablets collected from different manufacturers were detected.

#### 4.2.2 Simultaneous Measurements of Content of Seven Components in Ginkgo Biloba Tablets by UPLC-MS Method

##### Preparation of Test Sample Solution

Quantitative (one tablet for samples numbered A1-A6 and two tablets for samples numbered B1-B4) was accurately weighed after being crushed and added into 5.0mL methylene chloride to dissolve. The sample bottle was sealed and soaked for eight hours and ultrasonically dispersed after the solvent fully infiltrated the tablet carrier. During the ultrasonic process, the solution was kept at a temperature below thirty degrees celsius by pausing and adding ice cubes. After the effective components in the sample were completely dissolved, the supernatant was centrifuged and filtered. Afterwards, 1ml was measured and diluted with acetonitrile for later use.

##### Preparation of Standard Solution

The reference substances including ginkgolide A, ginkgolide B, ginkgolide C and bilobalide, quercetin, isorhamnetin and appropriate amount of kaempferol were precisely weighed, respectively. Then, dichloromethane was added to prepare the reference substance solution of about 1mg· mL^-1^. With appropriate amount of each reference substance solution taken respectively, methylene chloride was added to prepare mixed reference substance solution with appropriate concentration. They were stirred up sufficiently and filtered using 0.45μm filter membrane for later use.

##### Preparation of Series of Concentration Standard Solution

0.5, 1.0, 2.0, 4.0 and 8.0mL of the above mixed reference solution were measured accurately and placed in 10mL volumetric flasks respectively. Acetonitrile was added, diluted to scale and stirred up sufficiently for later use.

##### Chromatographic Conditions

Waters Xbridge C18 (4.6×150mm,3.5um) column was used, mobile phase A was acetonitrile and mobile phase B was water (containing 0.10% formic acid). It is gradient eluted (0 to 2 min, 0%A→5%A; 2 to 4min, 5%A→95%A, 4 to 30min, 95%A). Prior to each injection, the mobile phase A-B (50:50) pre-equilibrium was applied for a period of 5min, the flow rate was 1.5mL·min^-1^, the column temperature was 30°C and the injection volume was 10μL.

##### Mass Spectrometry Conditions

Negative ion mode monitoring was carried out with Electron Spray Ionization. Quantitative mode was adopted. Scanning range m/z was 100∼1400,Capillary voltage was 3.8kv. Spray pressure was 60psi and ion source temperature was 650°C. The interface was heated and nitrogen was introduced throughout the process.

##### Determination Method

10μL of test sample solution was accurately measured. According to “chromatographic conditions” and “mass spectrometry conditions”, LC-MS was applied for determination.

## 5. Conclusions

In this study, a total of 7 components in ginkgo biloba tablets were determined simultaneously by UPLC-MS. Methodological investigation revealed that this method was capable of determining the content of ginkgo flavonoids and ginkgolides in ginkgo biloba tablets and that of ginkgo flavonoids and ginkgolides in ginkgo biloba leaves. This article analyzed the limitations and shortcomings of the experiment. First of all, ginkgo biloba is an extract in traditional Chinese medicine, which is made use of widely in China, Japan, South Korea, Korea and Southeast Asia. However, it remains rarely used in other countries and regions. Therefore, the retrieval of relevant literature is subjected to certain limitations. Secondly, due to the influence exerted by the areas of production, the processing of the original medicinal materials of ginkgo biloba tablets of each batch, the processing techniques and the content of 7 effective components in ginkgo biloba tablets are different to some extent. However, the results of the sample for the test showed that the quantity of 7 active ingredients in Ginkgo biloba tablets was appropriate to the standard. This method provided a reference and basis for the quality control of ginkgo biloba tablets, in addition to laying a foundation for the further pharmacokinetic study of ginkgo biloba tablets in the future.

## Author Contributions

For research articles with four authors. The study was designed by LM and SSW. The experiments,as well as data analysis were conducted by LM,JZ and WRJ.JZ contributed analysis tools;The results were interpreted by LM and SSW. All the authors have made critical revision to the important knowledge content.

## Acknowledgments

We appreciate all the reviewers for their suggestions.

## Abbreviations

The following abbreviations are used in this manuscript:

KAk: aempferol
ISO: isorhamnetin
GA: ginkgolide A
GB: ginkgolide B
GC: ginkgolide C
BB: bilobalide
ESI: Electron Spray Ionization
UPLC-MS: ultra-high performance liquid chromatography-mass spectrometer
HPLC-UV: high performance liquid chromatography-ultraviolet
HPLC-ELSD: high performance liquid chromatography-evaporative light-scattering detector
LC-M: Shigh performance liquid chromatography-mass spectrometer
RSD: Relative Standard Deviation

## References

He, W.Y. Summary of studies on the selection and breeding of improved ginkgo varieties and high-yield cultivation techniques in China. Gansu Nongye 2013, 378, 92–94.

Sun, F.; Wang, L.; Yan, B.; Peng, X. Chemical components and pharmacological effects of ginkgo biloba extracts. Shandong Journal of Traditional Chinese Medicine 2014, 33, 221–223.

Zhang, H.M. The chemical constituents and pharmacological activities of natural medicine, ginkgo biloba. Journal of Capital Normal University(Natural Sciences Edition) 2014, 35, 41–46.

Liu, X.P.; Zang, H.C.; Yu, H.L. Research progress and application prospect of ginkgo biloba extract. Journal of Pharmaceutical Research 2014, 33, 721–723.

Wang, G.Y.; Zhu, J.J.; Lou, F.C. Chemical constituents from the exopleura of Ginkgo biloba and inhibition test of total ginkgolic acids against phytopathogenic fungi. Journal of China Pharmaceutical University 2014, 45, 170–174.

Tian, Q.Y.; Gong, L.L. Research progress in ginkgo terpene lactones. Central South Pharmacy 2016, 14, 838–841.

Zhang, P.F.; Liao, L.J.; Deng, Z.; Tan, Y.P. Research progress of pharmacological effects and clinical application of ginkgo biloba extract. Liaoning Journal of Traditional Chinese Medicine 2017, 44, 426–429.

Bao, Y.R.; Zhang, H.R.; Liu, L.T. Optimization of extraction process of flavonoids from ginkgo biloba leaves and antioxidant activity. Journal of Henan University of Technology(Natural Science Edition) 2016, 37, 96–99.

Xu, X.G.; Yang, D.B.; Yu, R.C. Study on determination method of flavonoids in ginkgo leaves Shandong Pharmaceutical Industry 1999, 19, 20–21.

She, J.H.; Liu, Z.L. Analysis of flavonoid extracts from ginkgo leaves (ginkgo biloba) by SFE-HPLC. Chinses Traditional and Herbal Drugs 2000, 31, 101–103.

Bi, Y.M.; Sun, G.X.; Yu, X.M. Determination of ginkgolides in ginkgo leaf extract and dipyridamole injection by HPLC-ELSD. Central South Pharmacy 2007, 5, 124–127.

Yang, Y.F.; Xia, Y.Y. Progress on determination methods of terpene lactones in ginkgo leaves and their preparations. Xian Dai Yao Wu Yu Lin Chuang 2000, 15, 61–66.

Lv, F.S.; Chen, W.; Feng, F.; Zhang, Z.X. The contents of terpenelactones in the ginkgo injection. Journal of China Pharmaceutical University 2001, 32, 34–36.

Liu, H.; Huang, S.W.; Guo, Y.; Xiong, Y. Determination of flavonol glycosides and terpene lactones in Ginkgo biloba extract from different pharmaceutical factories by HPLC. China Journal of Traditional Chinese Medicine and Pharmacy 2015, 30, 598–601.

Gong, L.L.; Tian, J.Z. Determination of terpene lactones in ginkgo biloba extract by HPLC-ELSD. Journal of Shandong University of Traditional Chinese Medicine 2013, 37, 69–70.

Ou, Q.; Zhang, L.Y.; Qian, Y.X.; Kang, J.C.; He, J. Determination of flavonoids and lactone in shuxuening injection by HPLC. Journal of Guizhou University (Natural Science) 2010, 27, 34–37.

StatePharmacopoeiaCommittee. Pharmacopoeia of the People’s Republic of China (Part I) Beijing: China Pharmaceutical Science and Technology Association 2015, 1491–1492.

Qiao, S.; Shi, R.; Jiang, W.J.; Shu Jie, L.U.; Wang, Q.; Zhang, L.T. Simultaneous determination of six components in zixinyin oral liquid by LC-MS/MS. Chinese Traditional Patent Medicine 2011, 33, 456–460.

Liang, W.L.; Xie, D.W.; Ding, G.; Xu, D.H.; Sun, Y.C.; Lian, Y.P.; Li, Y.J.; Xiao, W. Quantitative and qualitative evalution on tablets of ginkgo biloba leaves using fingerprint and LC-MS analysis. China Journal of Chinese Materia Medica 2015, 40, 1738–1743.

Li, L.F.; Sun, G.X. Quantified fingerprints of ginkgo tablet by HPLC and content determination of four components. Chinese Journal of Experiment Traditional Medical Formulae 2014, 20, 52–57.

Tong, R.F.; Lin, H.; Li, W.C.; Zhang, N. Determination of ginkgolid A, B, and C, and bilobalide from Yinxingye Tablets by HPLC-MS method. Drugs & Clinic 2017, 32, 386–389.

Ren, Y.P.; Chang, L.; Cao, L.; Zhi, X.R.; Zhang, L.T. Simultaneous determination of terpene lactones and favonoid aglycones in Shuxuening injection by HPLC-MS. Chinese Journal of Pharmaceutical Analysis 2013, 33, 220–224.

Xu, H.J.; Ji, X.; Chen, Z.; Lu, S.J.; Zhang, L.T. Simultaneous determination of seven coumarins in qingxuan tablets by HPLC-MS. Chinese Pharmaceutical Journal 2011, 46, 1526–1529.

Ganzera, M.; Sturm, S. Recent advances on HPLC/MS in medicinal plant analysis—an update covering 2011-2016. Journal of Pharmaceutical and Biomedical Analysis 2017, 147, 211–233.

